# Exogenous Thyroxine Increases Cardiac Nrf2-TRX Associated with a Reduction in Oxidative Injury and Inflammation in Insulin Resistant OLETF Rats

**DOI:** 10.1101/2025.04.16.649239

**Authors:** Dora A. Mendez, Jenifer Hernández García, José G. Soñanez-Organis, Marisol Hernández Garcia, Guillermo Vazquez-Anaya, Akira Nishiyama, José Pablo Vázquez-Medina, Rudy M. Ortiz

## Abstract

Cardiovascular disease (CVD) is the leading cause of death among individuals with Type II diabetes (T2D), affecting approximately 30 million people in the United States. During insulin resistance, the heart undergoes a metabolic shift, leading to increased reactive oxygen species (ROS) generation, lipotoxicity, and mitochondrial dysfunction, ultimately contributing to cardiovascular dysfunction. The effects of thyroid hormones (THs) on redox biology and oxidative stress remain inconclusive, necessitating further investigation. In this study, insulin-resistant Otsuka Long Evans Tokushima Fatty (OLETF) rats were used to assess the impact of exogenous thyroxine (exoT4) on NADPH oxidases (NOX) and antioxidant defenses in the heart. Rats were assigned to four groups: **(1)** lean control, Long Evans Tokushima Otsuka (LETO; n=6), **(2)** LETO + T4 (8 μg/100g BM/day for 5 weeks; n=7), **(3)** untreated OLETF (n=6), and **(4)** OLETF + T4 (n=7). NOX4 mRNA expression was two-fold greater in OLETF rats compared to LETO. T4 treatment increased NOX4 protein abundance by 56% in OLETF. Additionally, T4 normalized lipid peroxidation (4-hydroxynonenal) and tumor necrosis factor-α (TNF-α) levels while increasing nuclear factor erythroid 2-related factor 2 (Nrf2) mRNA expression by 158% compared to LETO and enhancing nuclear Nrf2 protein expression by 45% compared to untreated OLETF. Thioredoxin (TRX) expression, suppressed in OLETF, was increased by 88% following T4 treatment. ExoT4 increased mitofusin 2 (Mfn2) protein abundance in OLETF by 49% compared to LETO. These findings suggest that thyroid hormone treatment may have cardioprotective effects mediated by Nrf2 in the heart during metabolic syndrome (MetS).

## Introduction

Cardiovascular disease (CVD) continues to be the leading cause of death globally (1) costing the United States about $239.9 billion each year (2). CVD can be a primary condition or a secondary consequence of various other disorders, including diabetes, hypertension, metabolic syndrome, and obesity. According to the International Diabetes Federation, currently, 537 million people live with diabetes (3). Diabetes, a chronic and metabolic disease, is characterized by insulin resistance and high blood glucose, which over time, increase the risk of developing heart disease (4). These factors are concurrent with those seen in Metabolic Syndrome (MetS), which express clusters of interconnected risk factors, including abdominal obesity, impaired glucose tolerance with or without insulin resistance, high triglyceride levels, low HDL cholesterol, and/or hypertension (5). Insulin resistance is a key underlying characteristic of the pathophysiology of both type 2 diabetes and MetS. Animal models of MetS demonstrate cardiac metabolic inflexibility characterized by impaired glucose uptake and metabolism contributing to cardiac mitochondrial impairment (6). This metabolic shift causes an increase in fatty acid metabolism that further suppresses cardiac glucose metabolism (7). Consequences of altered fatty acid metabolism include myocardial lipotoxicity, mitochondrial dysfunction, and increased oxidative stress, all contributing to cardiac dysfunction (8).

Some major sources of reactive oxygen species (ROS) in the heart include but are not limited to, NADPH oxidases (NOX) and mitochondria (9). Under normal conditions, electrophiles are essential for normal signal transduction and maintaining cellular homeostasis and other biological functions. Oxidative stress arises when the production of oxygen radicals exceeds the antioxidant capacity and promotes the oxidation of cellular compounds, impairing their functions. Excessive ROS generation poses a threat to biological systems due to their ability to interact with various macromolecules, including DNA lipids, and proteins (10). Increased free fatty acids (FFA) lead to mitochondrial dysfunction, causing uncoupling of oxidative phosphorylation and elevated superoxide O_2_•– production, which induce oxidative injury (11). NOX2 and NOX4 are the predominant NADPH oxidase isoforms expressed in cardiac muscle. NOX2 resides in the plasma membrane and requires the recruitment of other cytosolic subunits (p47, p40, p67, p21RAC1) for activation (12) and generates O_2_•– when activated. NOX4 activation requires no cytosolic regulatory subunits beyond its membrane conjugate, p22phox, localizes to intracellular membranes, and generates primarily hydrogen peroxide (H_2_O_2_) and to a lesser extent, O_2_•– (13). Elevations in cardiac NOX4 have been implicated in a hormetic response in rats with MetS (14), offering CV protection against cardiac remodeling. In addition, NOX4-derived H_2_O_2_ may upregulate the transcription factor, nuclear factor erythroid 2-related factor 2 (Nrf2), which is responsible for upregulating antioxidant genes (14, 15).

Recent studies suggest that thyroid hormone treatment during cardiovascular complications has potential cardioprotective effects. Exogenous TH (T3 or T4) treatments protect the heart from oxidative and/or inflammatory injury following induced acute myocardial infarct (AMI) (16). Other studies revealed TH treatments rescued mitochondria function, attenuated cardiac remodeling, and counteracted diabetes-induced pathological remodeling, demonstrating their potential to ameliorate CV defects regardless of etiology (17-19). Previous work suggests that thyroid hormones (THs) can modulate the redox balance of cardiac cells under various stress conditions through multiple mechanisms: **(1)** membrane-initiated pathways for the protein kinase B (AKT)/endothelial nitric oxide synthase (eNOS) axis, and **(2)** transcriptional regulation of Nrf2, peroxisome proliferator-activated receptor gamma coactivator 1-alpha (Pgc1α) and transcription factor A, mitochondrial (Tfam), which upregulate antioxidant enzymes such as superoxide dismutases (SOD), glutathione peroxidases (Gpx), and peroxiredoxins (PRDX) (20-23). However, contradicting data also exists showing that TH treatments are associated with or promote hepatic oxidative stress (24-26). Thus, the effects of TH on redox biology are incongruent and maybe tissue-specific, requiring further investigation.

Previously, we have shown that the Otsuka Long Evans Tokushima Fatty (OLETF) rat exhibits cardiac oxidative injury and suppressed antioxidant capacity compared to the lean, control strain, Long Evans Tokushima Otsuka (LETO) rat (14). However, the effects of an exogenous thyroid hormone treatment on redox biology in the heart under insulin-resistant conditions associated with MetS remain poorly understood. Here, we investigated the effects of exogenous thyroxine treatment on cardiac NOX expression, antioxidants, and mitochondrial function in OLETF rats, a model of MetS.

## Methods

All experimental procedures received approval from the Institutional Animal Care and Use Committees of both Kagawa Medical University, Japan, and the University of California, Merced, USA. This dataset extends a previous study conducted on the same animals, where we examined the impact of exogenous thyroxine on glucose intolerance (27). At the conclusion of the study, the hearts were quickly dissected, weighed, and snap-frozen in liquid nitrogen, then stored at −80°C for subsequent analysis. Sample integrity was confirmed through multiple qRT-PCR and Western blot analyses.

### Animals

Lean, male (265 ± 7 g) Long Evans Tokushima Otsuka (LETO) rats and obese, male (356 ± 4 g) Otsuka Long Evans Tokushima Fatty (OLETF) rats (Otsuka Pharmaceutical Co. Ltd., Tokushima, Japan), both 9 weeks old, were divided into the following groups (n = 6–7 per group): **(1)** untreated, control LETO, **(2)** LETO + T4 (8.0 μg/100 g BM/day for 5 weeks), **(3)** untreated OLETF, and **(4)** OLETF + T4 (8.0 μg/100 g BM/day for 5 weeks). The rats were housed in pairs in a specific pathogen-free facility at Kagawa Medical University, Japan. They were maintained at a controlled temperature of 23°C and 55% humidity, with a 12-hour light/dark cycle. All animals had ad libitum access to water and standard laboratory chow.

### T4 Administration

Osmotic minipumps (Alzet, model 2006, Durect Corp., Cupertino, CA), filled with L-thyroxine (T4) (Sigma-Aldrich, St. Louis, MO) dissolved in 6.5 mM NaOH and 50% propylene glycol, were implanted subcutaneously to deliver a set dose of 8.0 μg/100 g body mass per day for 5 weeks.

### Western Blots

A 20 mg sample of frozen heart tissue was processed using a two-step protocol to extract cytosolic and membrane fractions, as previously described (28). Briefly, the tissue was homogenized in 50 mM potassium phosphate buffer (Fisher Scientific, P290 and P285) containing 3% protease inhibitor cocktail (Sigma-Aldrich, P2714) and 3% phosphatase inhibitor cocktail (Thermo Scientific 78426). The homogenate was centrifuged at 15,000 × g to isolate the cytosolic fraction from the supernatant. The remaining pellet was then homogenized in 50 mM potassium phosphate buffer with 1% Triton X-100 (Millipore-Sigma, T8787) and the same inhibitor cocktails to extract the membrane fraction. Following a second centrifugation at 15,000 × g, the plasma membrane fraction was recovered from the supernatant. A 40 mg piece of frozen heart tissue was used to isolate mitochondrial and nuclear fractions as previously described (29). Following the subcellular extractions, protein concentrations were determined using the BCA protein assay (ThermoFisher, 23,221 and 23,224, Houston, TX, USA). Protein extracts (5-20 μg) were separated on 7.5–10% Tris-HCl SDS-PAGE gels. Before data collection, the purity of the cytosolic and membrane fractions was validated by testing against Na⁺/K⁺ ATPase (Abcam, ab76020, Boston, MA, USA) and α-tubulin (Abcam, ab52866, Boston, MA, USA), respectively. The purity of cytosolic, mitochondrial, and nuclear fractions was validated by testing against α-tubulin, voltage-dependent anion channels (VDAC) (Proteintech, 55259-1-AP, Rosemont, IL, USA), and histone-H3 (Proteintech, 17168-1-AP, Rosemont, IL, USA), respectively (Supplemental Fig 1). Membranes were incubated with primary antibodies against NADPH oxidase 2 (NOX2/gp91phox) (Abcam, ab310337, Boston, MA, USA), NADPH oxidase 4 (NOX4) (Proteintech, 14347-1-AP, Rosemont, IL, USA), nuclear factor (erythroid-derived 2)-like 2 (Nrf2) (Proteintech, 16396-1-AP, Rosemont, IL, USA), thioredoxin (TRX) (Abcam, ab26320, Boston, MA, USA), peroxiredoxin 6 (PRDX6) (Proteintech, 13585-1-AP, Rosemont, IL, USA), mitofusin 2 (Mfn2) (Proteintech, 12186-1-AP, Rosemont, IL, USA), PTEN-induced putative kinase 1 (PINK) (Proteintech, 23274-1-AP, Rosemont, IL, USA), parkin2 (Proteintech, 14060-1-AP, Rosemont, IL, USA), ATP-binding cassette, sub-family B (MDR/TAP), member 10 (ABCB10) (Proteintech, 14628-1-AP, Rosemont, IL, USA), and TNF-alpha (Proteintech, 17590-1-AP). Membranes were washed and incubated with secondary antibodies—either IR Dye 680RD donkey anti-rabbit or IR Dye 800CW donkey anti-rabbit (1:10,000 dilution). Blots were visualized using the Li-Cor Odyssey Infrared Imaging System and quantified with ImageJ (NIH, Bethesda, MD) and Image Studio Lite version 5.2 (Li-Cor). Uniform sample loading was confirmed by Ponceau staining, following a modified Nakamura method (30). Membranes were treated with Ponceau S solution (0.1% [w/v] Ponceau S in 5% [v/v] acetic acid) for 10 minutes and then rinsed with distilled water until distinct bands appeared. For mitochondrial proteins where loading required less than 5 µg/µL of total protein, VDAC served as the housekeeping protein to confirm uniform sample loading. Results are expressed as percent expression normalized to the LETO control.

### Cardiac enzyme activities, oxidative damage, and succinate content

Cardiac 4-hydroxynonenal (4-HNE) levels were quantified in crude extracts using commercially available EIA kits (Cell BioLabs, San Diego, CA). The activities of superoxide dismutase (SOD), catalase, and glutathione peroxidase (GPx) were assessed in crude extracts using commercially available kits (Cayman Chemical, Ann Arbor, MI). Succinate-coenzyme Q reductase (complex II; SDH) activity was determined in heart mitochondria following established protocols (31).

### Quantification of NOX2, NOX4, and NRF2 gene expression

Total RNA was isolated using TRIzol reagent (Invitrogen, Waltham, MA, USA). Its integrity was evaluated by 260/280 nm ratio absorbance and agarose gel electrophoresis. Genomic DNA was degraded using DNase I (Roche, Indianapolis, IN, USA), and cDNA was synthetized using 1 μg total RNA with the QuantiTect Reverse Transcription Kit (Qiagen) y oligo-dT (50 µM)). Quantitative PCR was performed for NOX2, NOX4, and NRF2, with primers designed by (113) (Table 1). Two PCR reactions for each cDNA were run using the Step-One Real Time PCR System (Applied Biosystems, Foster City, CA, USA) in a final volume of 15 μL with 60 ng of equivalent total RNA. After an initial denaturing step at 94 °C for 10 min, amplifications were performed for 40 cycles at 94 °C for 15 s and 63 °C for 1 min, and a final melting curve program increased 0.3 °C each 20 s from 60 to 95 °C. Negative controls were included for each gene. The mRNA concentration was obtained using standard curves from dilutions from 5E−4 to 5E−8 ng μL−1 of each PCR fragment and normalized to the expression of α-actin.

**Table 1.**
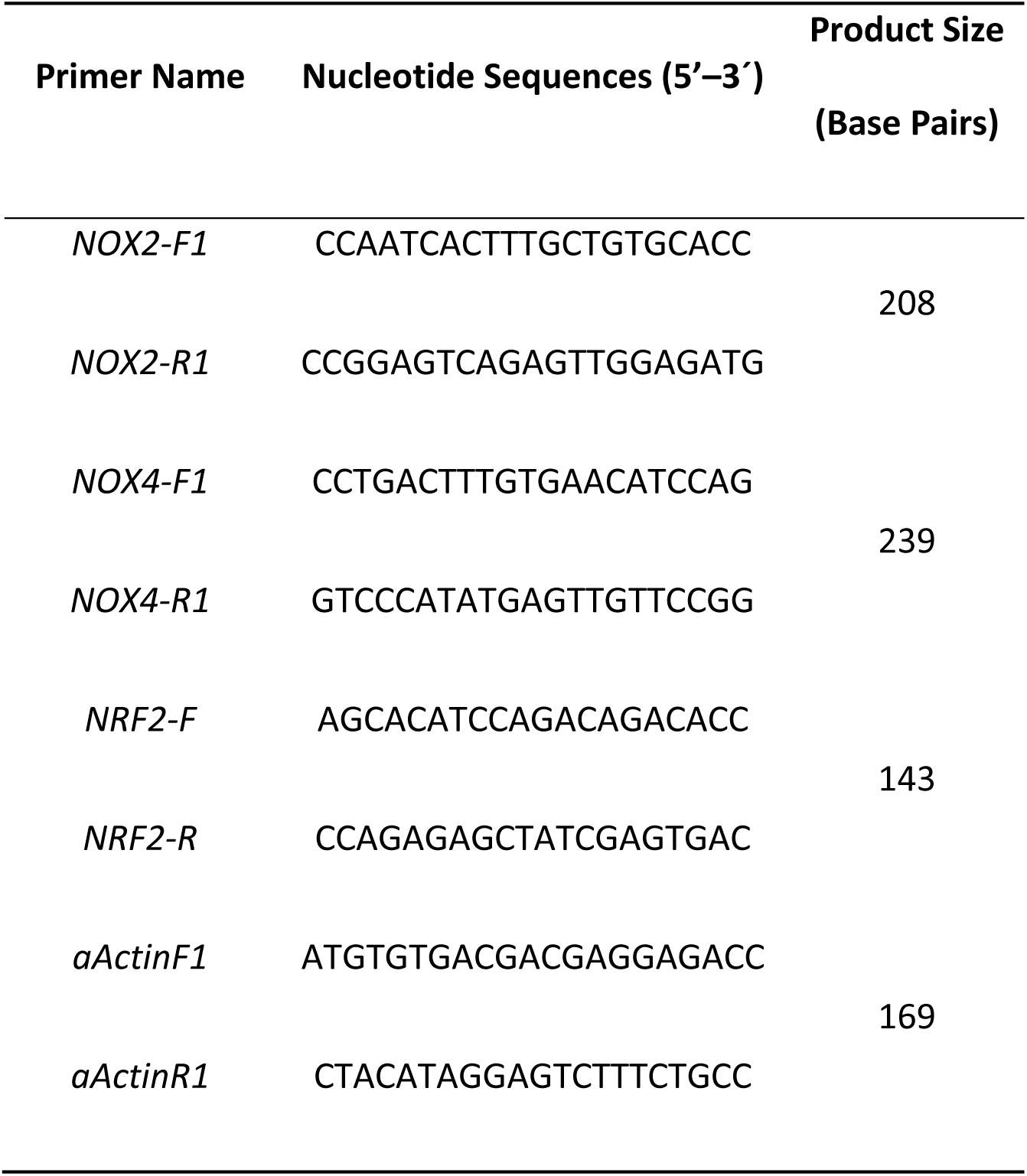
Primers used for the quantitative PCR.

### Statistics

Means (±SD) were analyzed using Two-way ANOVA with Tukey’s HSD post hoc test or an unpaired one-tailed t-test, with significance set at p < 0.05. The untreated LETO group served as the control for all comparisons to evaluate relative changes. Statistical analyses were conducted using GraphPad Prism 9.4.1 software (GraphPad Software).

## Results

### T4 increases mitochondrial NADPH Oxidase 4

NADPH oxidase gene and protein expression was measured to assess the effects of exogenous thyroxine on the potential for NOX-derived oxidant production and signaling in insulin-resistant hearts. Both treated and untreated OLETF groups showed higher NOX2 gene expression compared to LETO control (Fig 1A). However, these changes did not translate to parallel changes in protein abundance (Fig 1B). NOX4 mRNA was 2-fold greater in OLETF compared to LETO (Fig 1C). Both treated LETO and OLETF also exhibited higher NOX4 mRNA levels compared to LETO; however, treatment with T4 decreased NOX4 gene expression in OLETF compared to OLETF control (Fig 1C). Interestingly, NOX4 protein abundance in the mitochondrial fraction was reduced by 35% in OLETF compared to LETO. Treatment with T4 increased NOX4 protein abundance by 56% in OLETF (Fig 1D).

**Figure 1.**
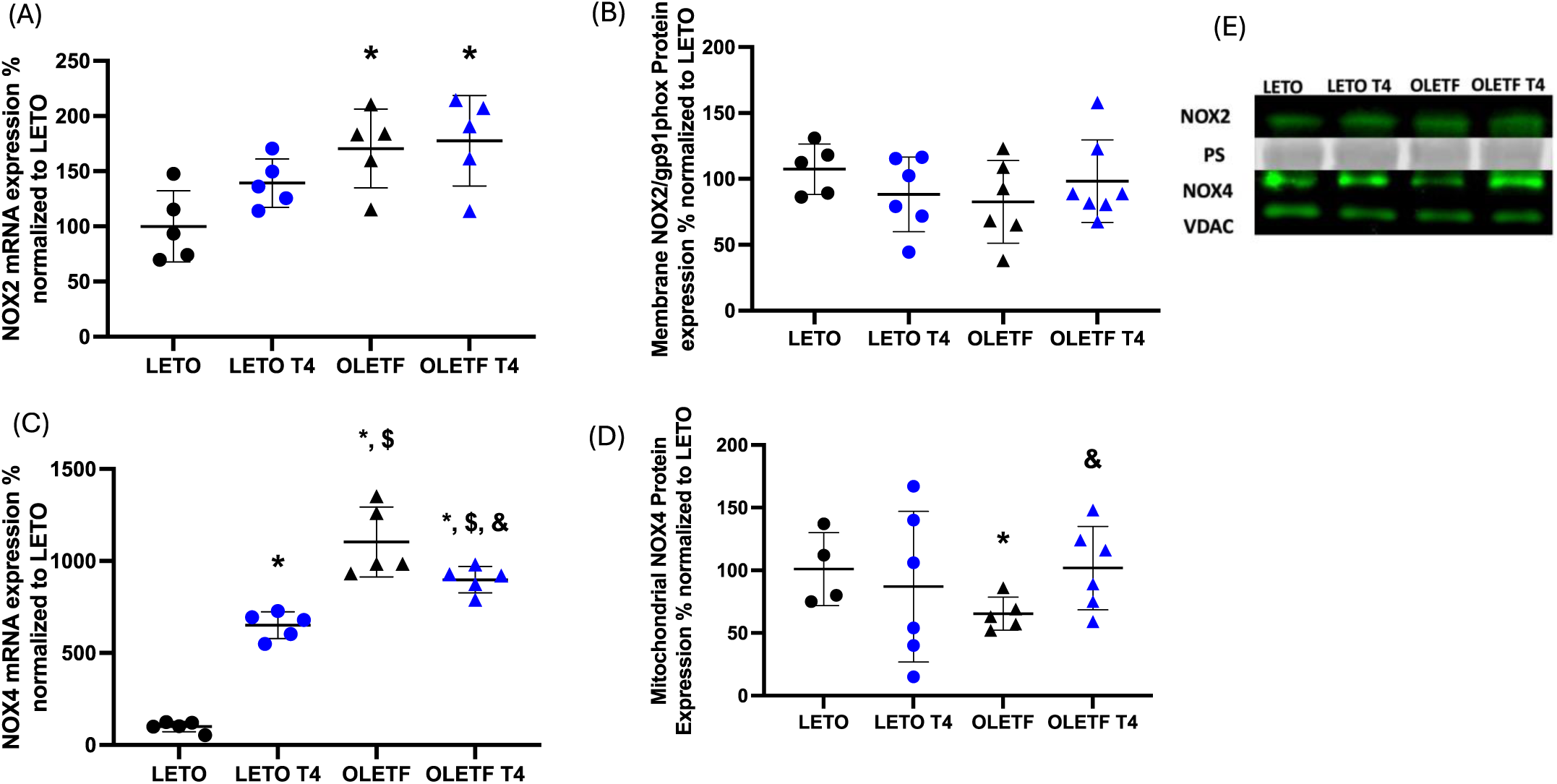
NADPH Oxidase gene and protein expression. Mean ± SD values for **(A)** NOX2 mRNA, **(B)** NOX2 protein expression in plasma membrane, **(C)** NOX4 mRNA, and **(D)** NOX4 protein expression in mitochondria. **(E)** Representative blots for NOX2 and NOX4 protein expressions of LETO, LETO + T4, OLETF, and OLETF + T4 rats (n = 6–7). PS = ponceau stain, * significantly different from LETO p < 0.05, ^&^ significantly different from OLETF p < 0.05, ^$^ significantly different from LETO + T4 p < 0.05, ^#^ significantly different from OLETF + T4 p < 0.05.

### T4 increases Nrf2 & TRX

To evaluate the effects of exoT4 on the redox-regulated cellular antioxidant defense in the hearts of obese, insulin-resistant rats, nuclear factor erythroid-2 related factor 2 (Nrf2) mRNA, nuclear protein expression, thioredoxin (TRX), and peroxiredoxin 6 (PRDX6) were quantified. Mean Nrf2 mRNA expression increased 73% in treated LETO and OLETF control compared to LETO. ExoT4 increased Nrf2 mRNA expression in OLETF by 158% compared to LETO (Fig 2A). Interestingly, Nrf2 protein abundance in the nucleus decreased 20% in LETO T4 and untreated OLETF compared to LETO. ExoT4 corrected nuclear Nrf2, increasing its expression by 45% compared to untreated OLETF (Fig 2B). TRX was suppressed in OLETF compared to LETO and exoT4 increased its protein expression by 88% compared to untreated OLETF (Fig 2C). Protein levels of PRDX6 increased 44% in untreated OLETF compared to LETO, and this elevation was not observed in the treated LETO and OLETF groups (Fig 2D).

**Figure 2.**
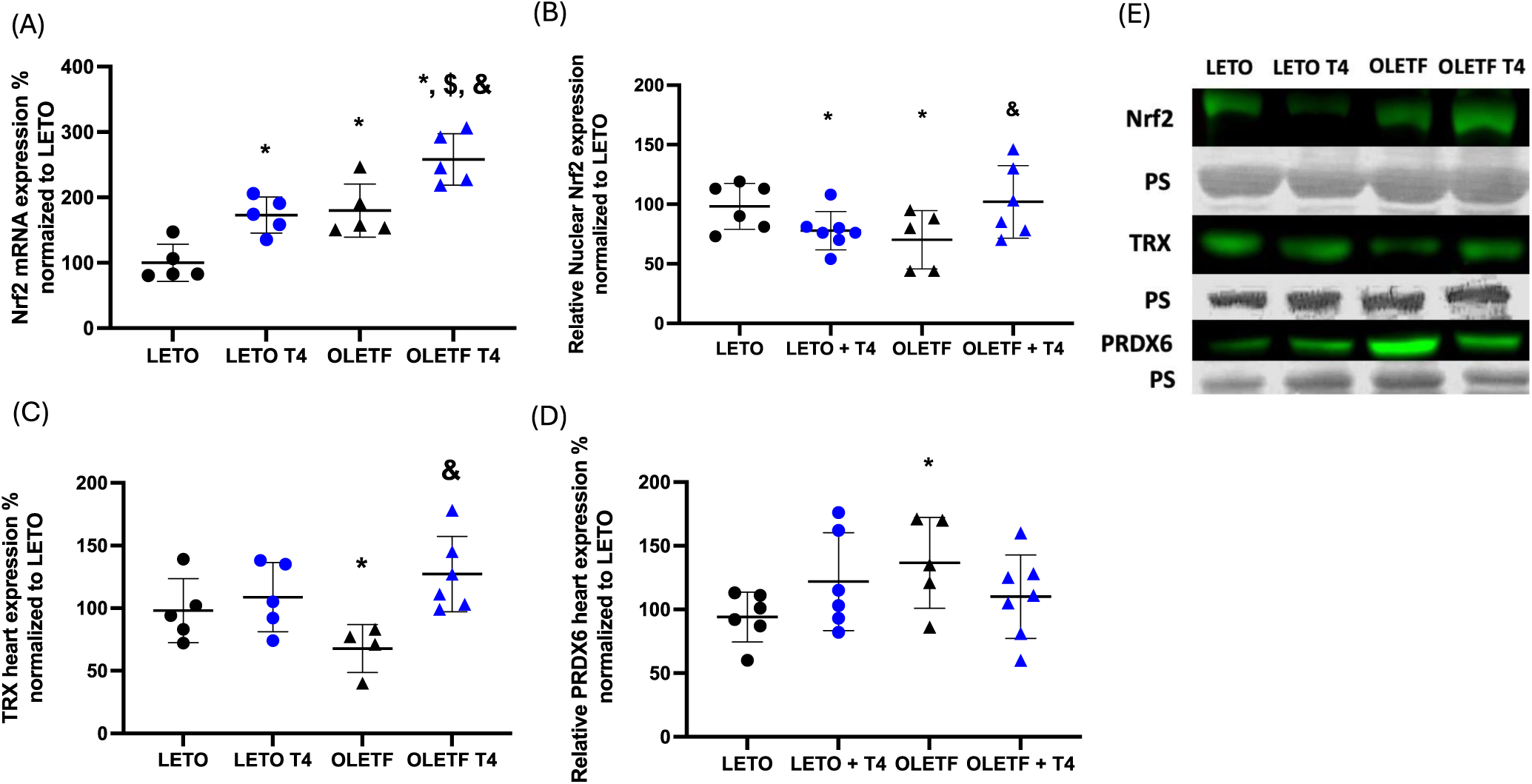
T4 increases Nrf2. Mean ± SD values for **(A)** Nrf2 mRNA, **(B)** Nrf2 protein expression in the nucleus, **(C)** thioredoxin (TRX) protein expression, and **(D)** peroxiredoxin 6 (PRDX6) protein expression. **(E)** Representative blots for the proteins shown in A-D of LETO, LETO + T4, OLETF, and OLETF + T4 rats (n = 6–7). PS = ponceau stain, * significantly different from LETO p < 0.05, ^&^ significantly different from OLETF p < 0.05, ^$^ significantly different from LETO + T4 p < 0.05, ^#^ significantly different from OLETF + T4 p < 0.05.

### Glutathione peroxidase & superoxide dismutase activities are suppressed in OLETF

The activities of catalase, glutathione peroxidase (GPx), and superoxide dismutase (SOD) were quantified to determine the impact of exoT4 on the heart’s antioxidant capacity in obese, insulin-resistant rats. Neither a strain nor treatment effect was detected for catalase activity (Fig 3A); however, a strain effect was detected for GPx and SOD (Fig 3B and 3C). Mean GPx and SOD activities were decreased by 33% and 44%, respectively, in untreated OLETF compared to LETO control and exoT4 generated an intermediary phenotype that trended toward normalization of activities for both (Fig 3B & Fig 3C).

**Figure 3.**
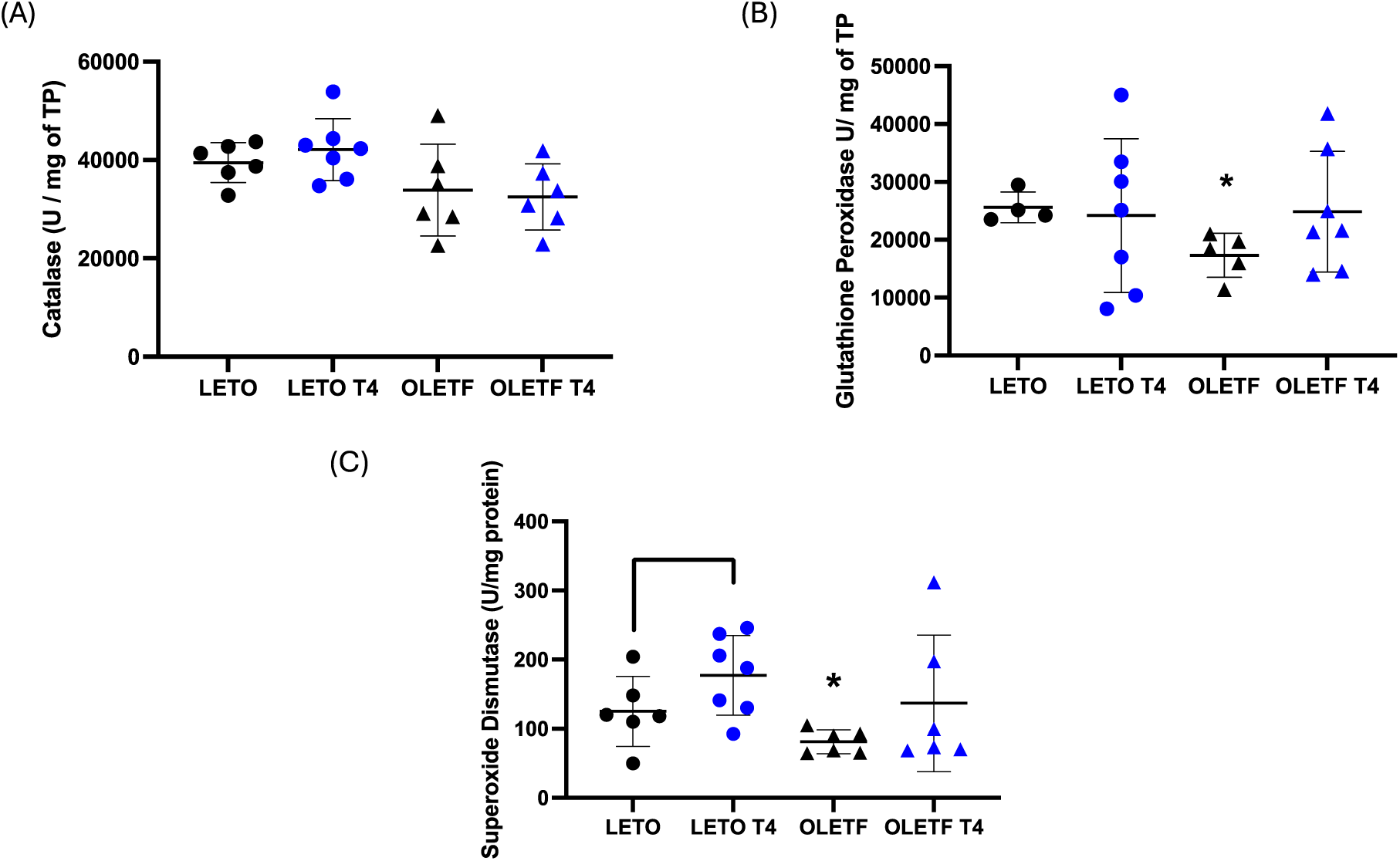
Strain effect in GPx and SOD. Mean ± SD values for activities of **(A)** catalase, **(B)** glutathione peroxidase (GPx), and **(C)** super oxide dismutase (SOD). * significantly different from LETO p < 0.05, ^&^ significantly different from OLETF p < 0.05, ^$^ significantly different from LETO + T4 p < 0.05, ^#^ significantly different from OLETF + T4 p < 0.05.

### T4 did not exacerbate cardiac 4-HNE and TNF-α

Cardiac 4-HNE levels and tumor necrosis factor-α (TNF-α) were measured to assess the effects of exoT4 treatment on activation of cardiac oxidative damage and inflammatory response in a model of insulin resistance. The non-treated OLETF group showed a 45% increase in cardiac 4-HNE levels compared to the LETO control strain (Fig 4A). There was no change observed in the treated LETO and OLETF compared to the LETO control strain (Fig 4A). The untreated OLETF group also expressed a 41% increase in TNF-α protein abundance compared to the LETO control strain (Fig 4B), no changes were observed in treated LETO and OLETF. ExoT4 generated an intermediary phenotype with trends toward improving these markers of injury and inflammation.

**Figure 4.**
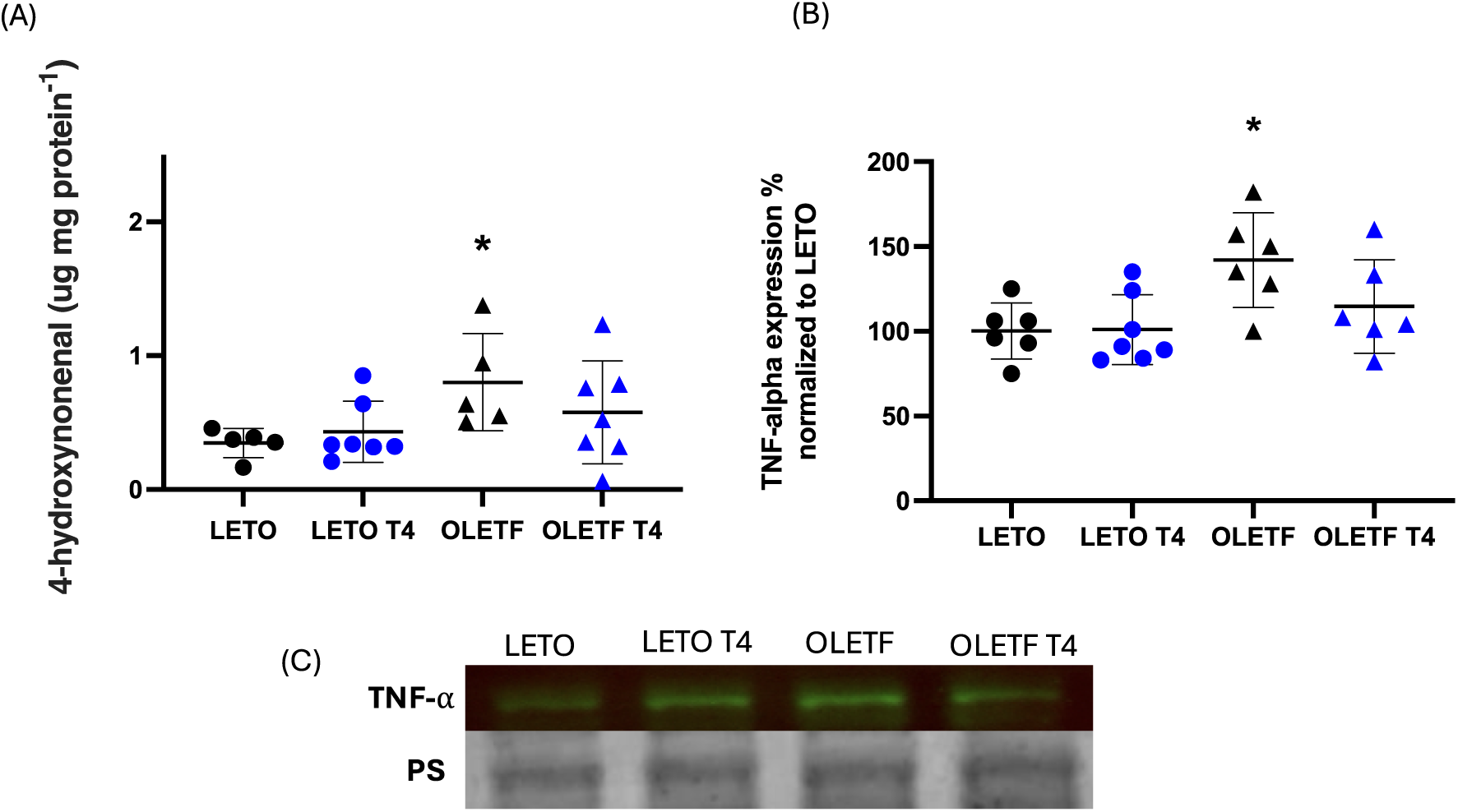
T4 partially reduced 4-HNE and TNF-α. Mean ± SD values for **(A)** cardiac 4-hydroxynonenal content, and **(B)** TNF-alpha protein expression. **(C)** Representative blot for TNF-alpha of LETO, LETO + T4, OLETF, and OLETF + T4 rats (n = 6–7). PS = ponceau stain, * significantly different from LETO p < 0.05, ^&^ significantly different from OLETF p < 0.05, ^$^ significantly different from LETO + T4 p < 0.05, ^#^ significantly different from OLETF + T4 p < 0.05.

### T4 increases mitofusin 2 & PINK1 protein expression

The expression of the mitochondrial proteins, mitofusin 2 (MFN2), ATP binding cassette 10 (ABC10), PTEN-induced putative kinase 1 (Pink1), and parkin, as well as the activity of succinate dehydrogenase (SDH) were measured to assess the contributions of exoT4 on mitochondrial homeostasis, function, and potential protection from oxidative injury in the hearts of insulin resistant OLETF rats. ExoT4 increased the mean protein expressions of Mfn2 and PINK1 in OLETF by 49% and 300%, respectively, compared to LETO, and increased PINK1 by 317% compared to untreated OLETF (Fig 5A & 5B). Mean parkin expression was decreased by 78% in untreated OLETF compared to LETO control and exoT4 did not alter the expression (Fig 5C). Similar to the behavior of PINK1 protein expression, exoT4 increased ABCB10 by 525% and 174% in OLETF compared to LETO and untreated OLETF, respectively (Fig 5D). Mean SDH activity decreased by 41% in OLETF compared with LETO; however, a treatment effect with exoT4 was not detected (Fig 5E).

**Figure 5.**
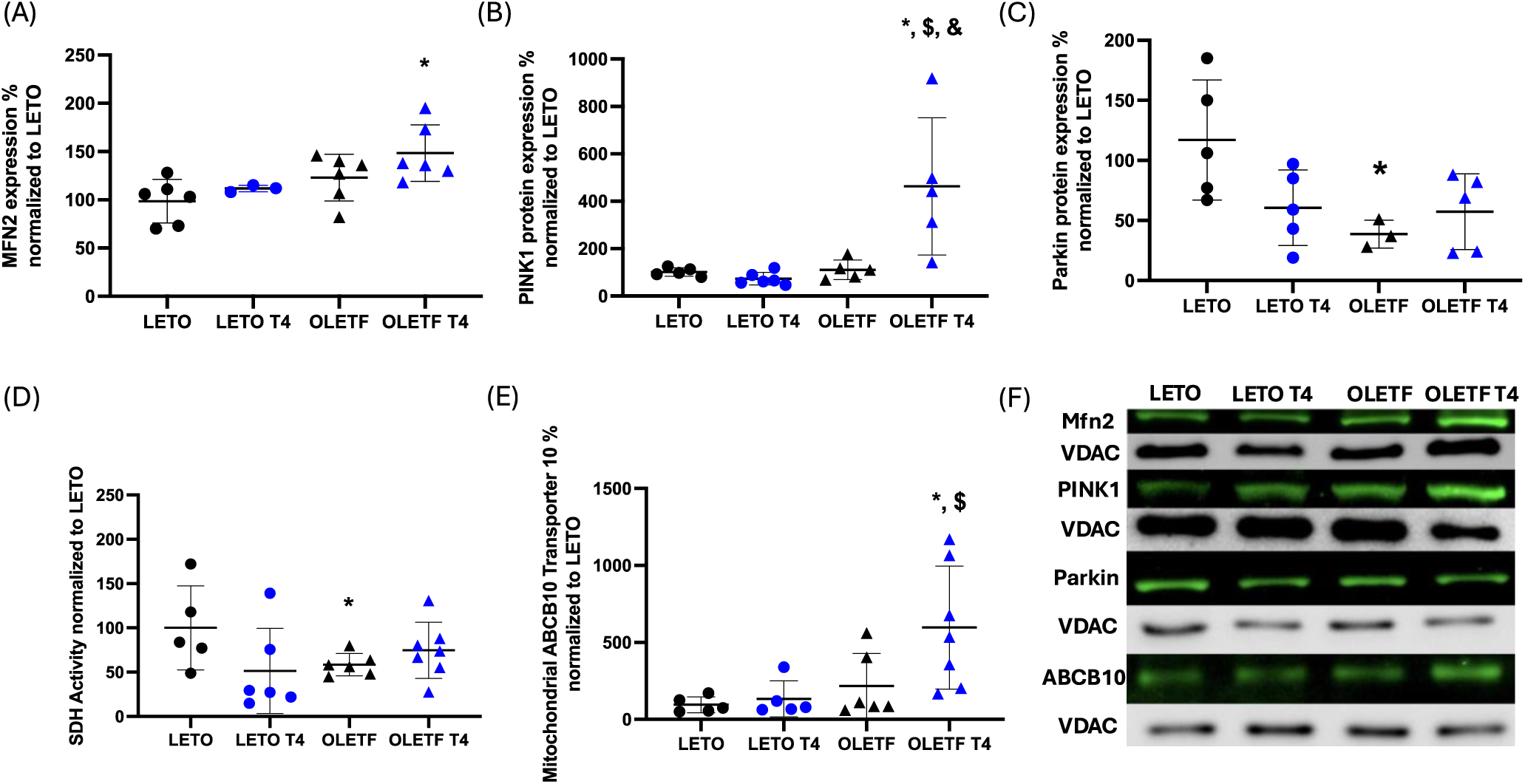
T4 increases Mfn2 and PINK1. Mean ± SD values for mitochondrial **(A)** mitofusin 2 (Mfn2) protein expression, **(B)** PTEN-induced putative kinase 1 (Pink1) protein expression, **(C)** Parkin protein expression, **(D)** ATP binding cassette 10 (ABC10) protein expression, and **(E)** succinate-coenzyme Q reductase (complex II; SDH) activity. **(F)** Representative blots for the proteins shown in A-D for LETO, LETO + T4, OLETF, and OLETF + T4 rats (n = 6–7). VDAC = mitochondrial voltage-dependent anion channels. * significantly different from LETO p < 0.05, ^&^ significantly different from OLETF p < 0.05, ^$^ significantly different from LETO + T4 p < 0.05, ^#^ significantly different from OLETF + T4 p < 0.05.

## Discussion

While data exists on the potential cardioprotective benefits of exogenous thyroid hormones (T3 or T4) during various conditions (8, 16, 53, 54), the data are incongruent, and the effect of exogenous T4 in the heart during MetS are not fully understood. In the present study, we demonstrated that exogenous T4 can increase cardiac antioxidant capacity and is associated with a trend towards amelioration of the strain-induced oxidative injury and inflammation and maintenance of mitochondrial homeostasis during MetS.

### NOXs

NADPH oxidases are a family of enzymes that produce ROS, specifically either superoxide (O_2_•–) or hydrogen peroxide (H_2_O_2_), as primary products during the catalytic metabolism of oxygen (33). At physiological levels, H_2_O_2_ participates in metabolic regulation by transducing cellular signals t_o_ aid cellular adaptation to environmental changes and stressors (34). However, the inappropriate elevation and/or activation of NOX proteins may promote oxidative injury. Several studies have demonstrated that an elevation and activation of cardiac NOX2 is associated with heart failure (12). In the present study, both treated and untreated OLETF groups exhibited elevated NOX2 mRNA, suggesting that the OLETF strain inherently displays increased NOX2 gene expression, which may contribute to the observed cardiac dysfunction in this model. This finding is consistent with what we have shown previously in this model (14). NOX4, which localizes to intracellular membranes and does not require cytosolic regulatory subunits aside from p22phox, has been shown to have both beneficial and detrimental effects (34). Elevated NOX4 in the endothelium enhances vasodilatation and reduces blood pressure and provides protection against chronic load-induced stress in mouse hearts by promoting angiogenesis (35, 36). In contrast, excessive NOX4 activity leads to endothelial dysfunction, vasoconstriction, hypertension, and a proinflammatory state, contributing to atherosclerosis and aneurysm development (37-40). The greatest expression in mean NOX4 mRNA was measured in the insulin-resistant OLETF group, suggesting that the phenotypic cluster factors observed in the OLETF strain, such as hypertension and insulin resistance, may be a consequence of NOX4. Interestingly, exogenous thyroxine increased the mRNA expression of NOX4 in both LETO and OLETF groups; however, these elevated levels in the OLETF group were still lower than the untreated OLETF, indicative of an intermediary phenotype that may have contributed to some of the benefits observed. Additionally, NOX4 protein levels remained high in OLETF T4 group. These data suggest that while thyroxine treatment can partially reduce NOX4 mRNA expression, it does not significantly lower NOX4 protein levels in insulin-resistant conditions, indicating post-transcriptional regulation or other compensatory mechanisms are contributing to the maintenance of NOX4 protein and likely a hormetic effect to counter the elevated NOX2 induced by the MetS condition.

Interestingly, for both NOX2 and NOX4, the increases in gene expression did not translate to parallel changes in protein expression. However, the regulation of NOX protein content is complex and a number of factors, including transcription factors such as NF-κB, molecules influencing mRNA stability, epigenetic processes such as DNA methylation, post-translational modification of histones, and the role of non-coding RNA can contribute the translation and/or post-translational modification of the protein (55). As noted previously, the highest expression of NOX4 mRNA was measured in the OLETF group, which had the lowest abundance of NOX4 protein, suggesting that in addition to the regulatory mechanisms mentioned above, protein degradation might be enhanced in OLETF, and thus, may have contributed to the measured reduction in NOX4 protein levels in OLETF. If so, and if the increased levels of NOX4 mRNA promote a hormetic response, then increased protein degradation of cardiac NOX4 would be detrimental. Additionally, the increased levels in the gene transcripts measured here may not have been sufficient to translate an increase in protein content. Nonetheless, the increased mRNA levels of NOX2 and NOX4 in the untreated OLETF group correlated well with the increased levels of cardiac 4-HNE an TNF-alpha in the same group suggesting that the measured increases in the transcripts were sufficient to promote oxidative injury and inflammation. Furthermore, the maintenance of elevated NOX4 protein in the presence of reduced gene expression in the exoT4 group suggests that T4 may induce mechanisms to increase the translational efficiency of this protein to help maintain a healthier phenotype and cardioprotection against the MetS-induced insults.

### Nrf2

Nrf2 is a transcription factor that regulates cellular redox balance by activating the transcription of Phase II genes, which are the majority of antioxidant enzyme genes, when it binds to its antioxidant response element (ARE) in the nucleus (43). In the present study, treatment with T4 significantly increased Nrf2 gene expression and protein abundance in the nucleus, suggesting that exogenous thyroxine has the potential to upregulate antioxidant genes, which may translate to upregulation of the associated enzymes and potentially protect tissues against oxidative injury. Studies have demonstrated that intracellular H_2_O_2_ can trigger the Nrf2-mediated antioxidant response (15), suggesting that elevation in NOX4 may have a hormetic effect to stimulate Nrf2 to protect against oxidative stress. The thioredoxin (TRX) system is a key antioxidant system which maintains a reducing environment by facilitating electron transfer from nicotinamide adenine dinucleotide phosphate (NADPH) through TRX reductase to TRX, which then reduces its target proteins using highly conserved thiol groups (44). Here, we show that TRX, a downstream target of Nrf2, is significantly reduced in OLETF, and T4 reversed this effect. In addition, we see a suppression in other critical antioxidants, such as GPx and SOD in the OLETF, demonstrating that antioxidant capacity is potentially downregulated during MetS conditions. Collectively, these data suggest that T4 may be inducing a cardioprotective effect by increasing NOX4, stimulating Nrf2 and upregulating TRX in the heart of these obese insulin-resistant animals. This phenomenon has been previously observed in tumor cells (45). PRDX6 is another antioxidant enzyme that reduces peroxides and helps in maintaining cellular redox balance (46). The increased PRDX6 levels observed in OLETF may be an adaptive response to the potential increases in NOX4-derived H_2_O_2_ and further indicate that this antioxidant defense system is likely responding to the heightened need to ameliorate elevated peroxides to reduce the potential of oxidative injury. The combination of increased PRDX6 and decreased TRX in OLETF may indicate an imbalance in redox signaling in the heart during MetS. While PRDX6 upregulation attempts to mitigate ROS levels, the concurrent reduction in TRX means that the overall antioxidant capacity is compromised.

### Mitochondrial Function

Mitofusin is a protein localized on the outer mitochondrial membrane that contributes to maintaining mitochondrial homeostasis by regulating the dynamic processes of mitochondrial fission and fusion while also facilitating intracellular signaling (47). PINK1 phosphorylates Mfn2, which subsequently recruits and binds parkin, acting as a mitochondrial receptor for parkin. Parkin then ubiquitinates Mfn2, marking it for proteasomal degradation. This sequence of interactions, known as the PINK1-Parkin-Mfn2 pathway, is essential for preserving mitochondrial morphology and function (52). In the present study, OLETF treated with T4 expressed high levels of Mfn2 and PINK1 in the heart, suggesting that PINK1 is activating Mfn2. Their increased expression suggests that thyroxine enhances mitochondrial quality control mechanisms in the hearts of OLETF rats, contributing to mitochondrial homeostasis. Enhanced mitochondrial function and homeostasis can reduce oxidative stress, indicating that T4 may have beneficial effects on mitochondrial health. Additionally, succinate dehydrogenase (SDH) is a mitochondrial enzyme that supports metabolic function by participating in the tricarboxylic acid (TCA) cycle and the electron transport chain (ETC). SDH activity was reduced in skeletal muscles of diabetic rat models, suggesting that diabetes and other metabolic disorders may disrupt mitochondrial and metabolic homeostasis through effects on TCA cycle (48). Similar to what we have shown previously (14), cardiac SDH activity was reduced in OLETF animals indicating that mitochondrial function is impaired. While exoT4 did not completely reverse this reduction, the treatment induced an intermediary phenotype that suggests that a longer duration or higher dose may have completely reversed the suppression in SDH activity. The observed preservation of SDH activity compared to LETO suggests that thyroxine may protect mitochondrial function by maintaining SDH activity and Mfn2.

Furthermore, ATP-binding cassette (ABC) transporters are a large family of proteins that utilize energy from ATP hydrolysis to transport a wide range of molecules against their concentration gradient. The ABCB10 isoform is a mitochondrial transporter involved in redox balance and protection against oxidation (49). In cardiac dysfunction after ischemia/reperfusion, inactivation of 1 allele of ABCB10 increased susceptibility to oxidative stress (50). In the present study, we demonstrate an increase in mitochondrial ABCB10 in the OLETF group treated with exogenous thyroxine. Because Nrf2 promotes ABCB10 expression (51), the observed increases in both Nrf2 and ABCB10 in exoT4 treated OLETF rats provide compelling evidence for a mechanistic link between Nrf2 and ABCB10. Collectively, these results support the hypothesis that T4 confers mitochondrial protection and improves redox homeostasis in MetS, which may have implications for therapeutic strategies targeting mitochondrial dysfunction in insulin resistance. Additionally, the association between subclinical and overt hypothyroidism and CVD, especially impaired CV functions (56) provides a basis for these implications. The regression of impaired CV functions in patients with thyroid hormone deficiency on L-T4 replacement therapy substantiates these implications and these results provide potential mechanisms for these benefits (56).

### Limitations

A key limitation of this study is the tissue-specificity of exogenous T4 administration. While our research primarily focused on the effects of T4 on cardiac tissue, it is essential to consider that T4 may have varying impacts on other tissues, such as the brain, liver, and kidneys. Existing studies provide insights into T4’s effects on the liver, where it has been shown to promote oxidative stress. This inconsistency underscores the necessity of a more comprehensive analysis of T4’s systemic effects. Recognizing these potential differential effects is crucial for accurately interpreting our findings and their broader implications. Future studies should investigate the effects of exogenous T4 across multiple tissues to provide a more holistic understanding of its therapeutic potential. Direct activation of some NOX enzymes is achieved by either phosphorylation-dependent pathways or subunit assembly. Therefore, measurement of NOX phosphorylation levels or degree of subunit assembly may have provided greater insight on their potential contribution to redox signaling here especially since we did not observe parallel changes in NOX gene and protein expressions. Future studies should incorporate more detailed studies of NOX enzyme regulation.

### Perspectives

The present study underscores the complex role of NOX4 in the insulin-resistant heart and its modulation by exogenous thyroxine. Interestingly, T4 treatment in OLETF slightly decreased NOX4 mRNA associated with a modest increase in mitochondrial NOX4 protein expression suggesting that enhanced thyroid hormone signaling improves the efficiency of NOX4 translation and/or protein stability to help maintain redox balance. The upregulation of Nrf2 and its downstream antioxidant targets, such as thioredoxin (TRX), in response to T4 treatment suggests a potential hormetic effect, where NOX4-derived H₂O₂ stimulates protective pathways rather than promoting oxidative damage. Thus, the hormetic effects go beyond the simulation of Nrf2-mediated phase II gene transcription but may also include the stimulation of mechanisms that promote the improvements in mitochondrial function, including increased ABCB10 expression, preservation of succinate dehydrogenase (SDH) activity, and enhanced mitofusin 2 (Mfn2) levels, all of which contribute to improved mitochondrial homeostasis in the insulin-resistant heart.

Additionally, the interplay between thyroid hormone signaling, Nrf2, and NOX4 presents an intriguing avenue for future research. While increased NOX4 activity has been implicated in pathological conditions such as hypertension and atherosclerosis, its role in metabolic disease may be more context-dependent, potentially serving as a compensatory mechanism to counteract oxidative stress. Further studies are needed to elucidate the precise regulatory mechanisms governing NOX4 expression and activity in the insulin-resistant heart and to determine whether targeting this pathway could offer novel therapeutic strategies for mitigating cardiovascular complications in MetS and T2D.

## Summary

This study highlights that insulin resistance-associated MetS exacerbates oxidative injury, inflammation, and impaired redox signaling, as evidenced by increased lipid peroxidation, elevated TNF-alpha, and the suppression of Nrf2 and related antioxidants. Our findings demonstrate that exogenous thyroxine has significant cardioprotective potential in insulin-resistant OLETF rats by enhancing antioxidant defenses and preserving mitochondrial homeostasis. The ability of thyroxine to upregulate Nrf2 and TRX, while maintaining SDH activity and increasing Mfn2 and ABCB10 levels suggests that enhanced thyroid hormone signaling mitigates oxidative stress in the heart by improving mitochondrial integrity in the context of MetS and T2D. These insights substantiate the therapeutic potential of L-T4 in addressing cardiovascular and metabolic complications associated with insulin resistance and MetS.

## CRediT authorship contribution statement

Dora A. Mendez: Writing – review & editing, Writing – original draft, Supervision, Project administration, Investigation, Data curation, Conceptualization. Jenifer Hernández García: Writing-review & editing, Investigation. José G. Sonanez-Organis: Writing – original draft, Investigation. Marisol Hernández García: Writing-review & editing, Investigation. Guillermo Vazquez-Anaya: Project administration. Akira Nishiyama: Supervision, Project administration, Conceptualization. José Pablo Vázquez-Medina: Conceptualization, Writing-review & editing. Rudy M. Ortiz: Writing – review & editing, Supervision, Project administration, Funding acquisition, Data curation, Conceptualization.

## Data availability

Data will be made available on request.

## Acknowledgments

We thank Dr. Maria Zoghbi for her insight and valuable feedback on ABCB10. Schematic diagram (Fig. 6) created with Biorender.com.

**Fig 6.**
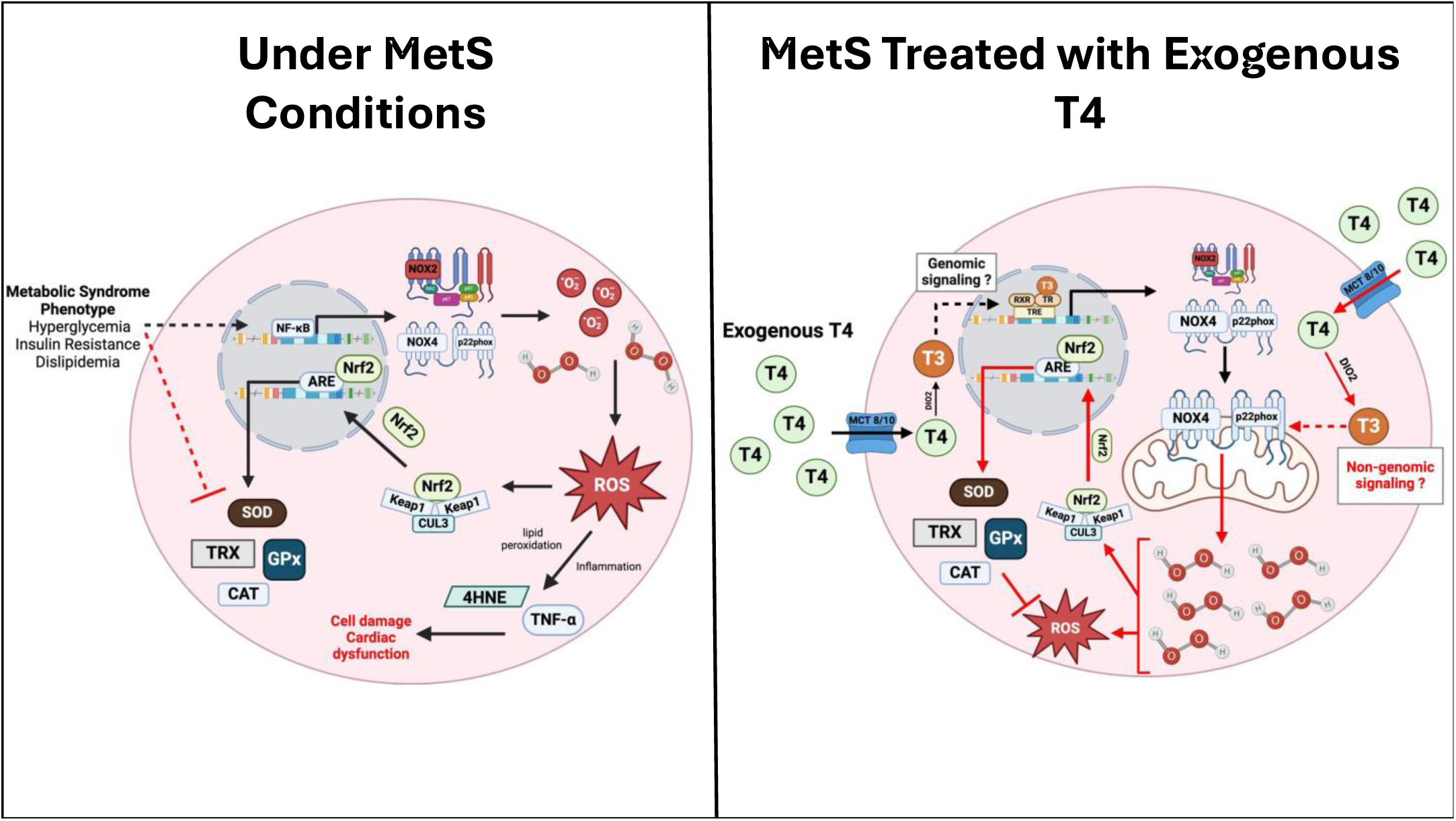
Schematic illustration of exogenous T4 on cardiac redox signaling. Left: Under metabolic syndrome (MetS) conditions. The MetS phenotype is indirectly (dashed black arrow) contributing to the increase in NOX2 and NOX4. Increases in NOX2 and NOX4 augment ROS generation. ROS triggers lipid peroxidation and inflammation, ultimately leading to cardiac cell damage and dysfunction. An increase in ROS generation also triggers the release of Nrf2 from Keap1, which then allows Nrf2 to migrate to the nucleus and activate downstream antioxidant signaling pathways. The MetS phenotype indirectly (red dashed inhibition line) suppresses antioxidants, contributing to the imbalance in redox and the potential for cardiac dysfunction. Right: Treatment with exogenous T4 in MetS conditions indirectly increases mitochondrial NOX4, the exact mechanism for this is not known. T4 could be increasing mitochondrial NOX4 via genomic (black arrows) or non-genomic signaling (red arrows). The increase in hydrogen peroxide released from NOX4 activates Nrf2 as a compensatory mechanism. Nrf2 then triggers downstream antioxidant signaling which ultimately reduce the imbalance in redox.

